# Dual-site TMS demonstrates causal functional connectivity between the left and right posterior temporal sulci during facial expression recognition

**DOI:** 10.1101/804971

**Authors:** Magdalena W. Sliwinska, Ryan Elson, David Pitcher

**Author notes:** **Corresponding author:** Magdalena W. Sliwinska. **Declaration of interest:** none.

## Abstract

**Background:** Neuroimaging studies suggest that facial expression recognition is processed in the bilateral posterior superior temporal sulcus (pSTS). Our recent repetitive transcranial magnetic stimulation (rTMS) study demonstrates that the bilateral pSTS is causally involved in expression recognition, although involvement of the right pSTS is greater than involvement of the left pSTS.

**Objective/Hypothesis:** In this study, we used a dual-site TMS to investigate whether the left pSTS is functionally connected to the right pSTS during expression recognition. We predicted that if this connection exists, simultaneous TMS disruption of the bilateral pSTS would impair expression recognition to a greater extent than unilateral stimulation of the right pSTS.

**Methods:** Participants attended two TMS sessions. In Session 1, participants performed an expression recognition task while rTMS was delivered to the face-sensitive right pSTS (experimental site), object-sensitive right lateral occipital complex (control site) or no rTMS was delivered (behavioural control). In Session 2, the same experimental design was used except that continuous theta-burst stimulation (cTBS) was delivered to the left pSTS immediately before behavioural testing. Session order was counter-balanced across participants.

**Results:** In Session 1, rTMS to the rpSTS impaired performance accuracy compared to the control conditions. Crucially in Session 2, the size of this impairment effect doubled after cTBS was delivered to the left pSTS.

**Conclusions:** Our results provide evidence for a causal functional connection between the left and right pSTS during expression recognition. In addition, this study further demonstrates the utility of the dual-site TMS for investigating causal functional links between brain regions.

## 1. Introduction

Humans recognise facial expressions using a distributed network of highly interacting brain regions (1, 2). One of these regions is located in the posterior superior temporal sulcus (pSTS). Prior neuroimaging studies of expression recognition (3–9) demonstrate that the right pSTS (rpSTS) shows a greater response to expressions than the left pSTS (lpSTS), leading to the suggestion that expression processing is right lateralized in the pSTS. Consequently, the role of the rpSTS in expression recognition has been extensively investigated while the role of the lpSTS remains relatively understudied.

We recently addressed this issue by using repetitive transcranial magnetic stimulation (rTMS) to demonstrate that both the right and left pSTS are important for accurate expression recognition (10). Crucially, while the impairment of an expression recognition task was greater in the rpSTS than in the lpSTS, stimulation to the lpSTS also impaired the task. This result shows that the contribution of the lpSTS should not be neglected when gaining a full understanding of expression recognition in the brain. One possible account of our result is that rTMS to the lpSTS reflects transient impairment of the lpSTS only, suggesting that the lpSTS contributes to the task independently from the rpSTS. However, it is equally possible that rTMS to the lpSTS was also impairing functional interactions between the left and right pSTS occurring via callosal connections. To further address this question, we conducted a dual-site TMS study on the bilateral pSTS.

Causal functional interactions between brain regions can be investigated using dual-site (so called condition-and-perturb) TMS (11). In this method, one brain region is conditioned with *off-line* TMS before participants perform a task and another region is then perturbed with *online* TMS during task performance. This method measures if task impairment caused by the perturbing TMS is greater following the conditioning TMS. Greater impairment of the perturbed region following the conditioning of another region demonstrates a functional interaction between the two regions, crucial for normal task performance. This method has been used in prior TMS studies to demonstrate functional connectivity between two regions within one hemisphere (11, 12) and across two hemispheres (13).

In the current study, functional magnetic resonance imaging (fMRI) was first used to identify the face-sensitive regions in the left and right pSTS and the object-sensitive region in the right lateral-occipital complex (rLO) for each participant. Then participants completed two TMS sessions (Session 1 and Session 2), session order was counter-balanced across participants. Both sessions involved three runs of a facial expression recognition task performed while perturbing rTMS was delivered either to the rpSTS (experimental run), right rLO (control site run) or no rTMS was delivered (control behavioural run). Perturbing rTMS was expected to impair the task when delivered to the rpSTS in contrast to the control conditions. The only difference between the two sessions was that in Session 2, conditioning continuous theta-burst (cTBS) stimulation was delivered to the lpSTS immediately before the behavioural testing commenced. We predicted that post-cTBS impairment of the lpSTS, lasting for at least 30 minutes (14, 15), would increase the impairment of the perturbing rTMS to the rpSTS, if the two regions are functionally connected.

## 2. Material and methods

### 2.1 Participants

Twenty-two right-handed participants were recruited in this study. Two participants found TMS uncomfortable and withdrew from the study while their data were discarded. All remaining participants (14 women and 6 men; aged between 19 and 25, mean: 20 years old, SD: 1.47 years old) were neurologically healthy with normal or corrected-to-normal vision.

Informed consent was obtained from all participants after the experimental procedures were explained. All participants were paid for their time. The study was approved by the York Neuroimaging Centre Research Ethics Committee at the University of York.

### 2.2 Experimental procedures

Each participant completed three sessions performed on different days. The first session involved the individual fMRI functional localisation to identify TMS target sites in every participant. The other two sessions involved TMS testing. The fMRI session lasted approximately 40 min while each TMS session lasted approximately 1 h.

### 2.3 Individual fMRI functional localisation

#### 2.3.1 Stimuli

Stimuli consisted of movie clips presenting moving faces or objects. Each movie clip lasted 3 sec and presented only one face or object. There were 60 movie clips for each stimulus category in which distinct faces or objects appeared multiple times. Moving faces were used to maximise the chance of finding face-sensitive areas in pSTS as this region was shown to respond stronger to dynamic stimuli than to the static stimuli, while activations for both types of stimuli spatially overlapped (16). These stimuli have also been used for localizing TMS target sites in our prior studies of the pSTS (10, 17, 18).

#### 2.3.2 Procedure

Functional localisation data were acquired over 2 block-design runs during which participants watched movie clips of dynamic faces or objects. Each run consisted of 10 blocks, 5 blocks per stimulus category. Within each block, 6 videos of either different faces or different objects were presented. Each block lasted 18 sec which made each run last 234 sec. Each run also contained three 18 sec rest blocks which occurred at the beginning, middle and end of the run. During the rest blocks, a series of six uniform colour fields were presented for 3 sec each. The order of stimulus category blocks in each run was palindromic (e.g., rest, faces, objects, faces, objects, faces, objects, rest, object, faces, object, faces, object, faces, rest) and randomised across runs. During this session, a structural brain scan was also acquired to anatomically localise the functional data for each participant.

#### 2.3.3 Data collection

Imaging data were collected using a 3T Siemens Magnetom Prisma MRI scanner (Siemens Healthcare, Erlangen, Germany) at the York Neuroimaging Centre. Functional localisation images from the whole brain were acquired using a 20-channel phased array head coil tuned to 123.3 MHz and a gradient-echo EPI sequence (60 interleaved slices; repetition time (TR) = 2000 msec; echo time (TE) = 30 msec; flip angle = 80°; voxel size = 3 × 3 × 3 mm; matrix size = 80 × 80; field of view (FOV) = 240 × 240 mm; total number of volumes per run = 117). Slices were aligned with the anterior to posterior commissure. Structural images were acquired using a high-resolution T1-weighted MPRAGE sequence (176 interleaved slices; repetition time (TR) = 2300 msec; echo time (TE) = 2.26 msec; flip angle = 8°; voxel size = 1 × 1 × 1 mm; matrix size = 256 × 256; field of view (FOV) = 256 × 256 mm).

#### 2.3.4 Data analysis

Functional localisation data were analysed for each participant using fMRI Expert Analysis Tool (FEAT) included in the FMRIB (v6.0) Software Library (www.fmrib.ox.ac.uk/fsl). In the first-level analysis, as part of the pre-statistical processing, single participant functional localisation images underwent extraction of non-brain structures using the Brain Extraction Tool (BET). In addition, interleaved slice timing correction, MCFLIRT motion correction, spatial smoothing using a 5mm full-width half-maximum Gaussian kernel, high-pass temporal filtering, and pre-whitening were applied to the data. To compute participant-specific patterns of activation, the pre-processed functional images were entered into a general linear model (GLM) with two independent predictors that correspond to the two stimulus categories (i.e., faces and objects). The model was convolved using a double-gamma hemodynamic response function (HRF) to generate the main regressors and temporal derivatives for each condition were included. Face-sensitive areas in the right and left pSTS were identified using a contrast of faces greater than objects. Object-sensitive areas in the rLO were identified using a contrast of objects greater than faces. First-level functional results were registered to the anatomical scan using a 6 degree-of-freedom affine registration. All analyses were conducted at the whole-brain level and voxel-level thresholding was set to p = 0.05. In the higher-level analyses, the first-level results for two runs of the functional localiser were averaged using fixed-effects and single group average model.

### 2.4 TMS study

#### 2.4.1 Stimuli

The stimuli in the expression recognition task comprised of 36 static images taken from Ekman and Friesen (19). Each image presented a face expressing an emotion. In total, faces of six female models (C, MF, MO, NR, SW, PF) were used and each model expressed six different emotions: happy, sad, surprise, fear, disgust, and anger. Each image was cropped with the same contour to cover the hair and neck of the models. Within each trial, the identity of the two faces was always different and within each run, the six expressions were presented an equal number of times.

#### 2.4.2 Procedure

TMS behavioural data were acquired over two sessions during which participants performed a computer-based expression recognition task. This task has been used in our previous studies and proved to be robustly affected by TMS to the right and left pSTS (10, 20, 21). During each trial (see Figure 1A), participants focused on the expressions of two people and were asked to judge whether their expressions were the same or not. The participants saw two images of faces presented sequentially for 500 msec each. The images were preceded by a fixation cross displayed for 2500 msec and separated by a fixation cross displayed for 1000 msec. Each trial ended with a blank white screen that was displayed until a response was provided.

**Figure 1:**
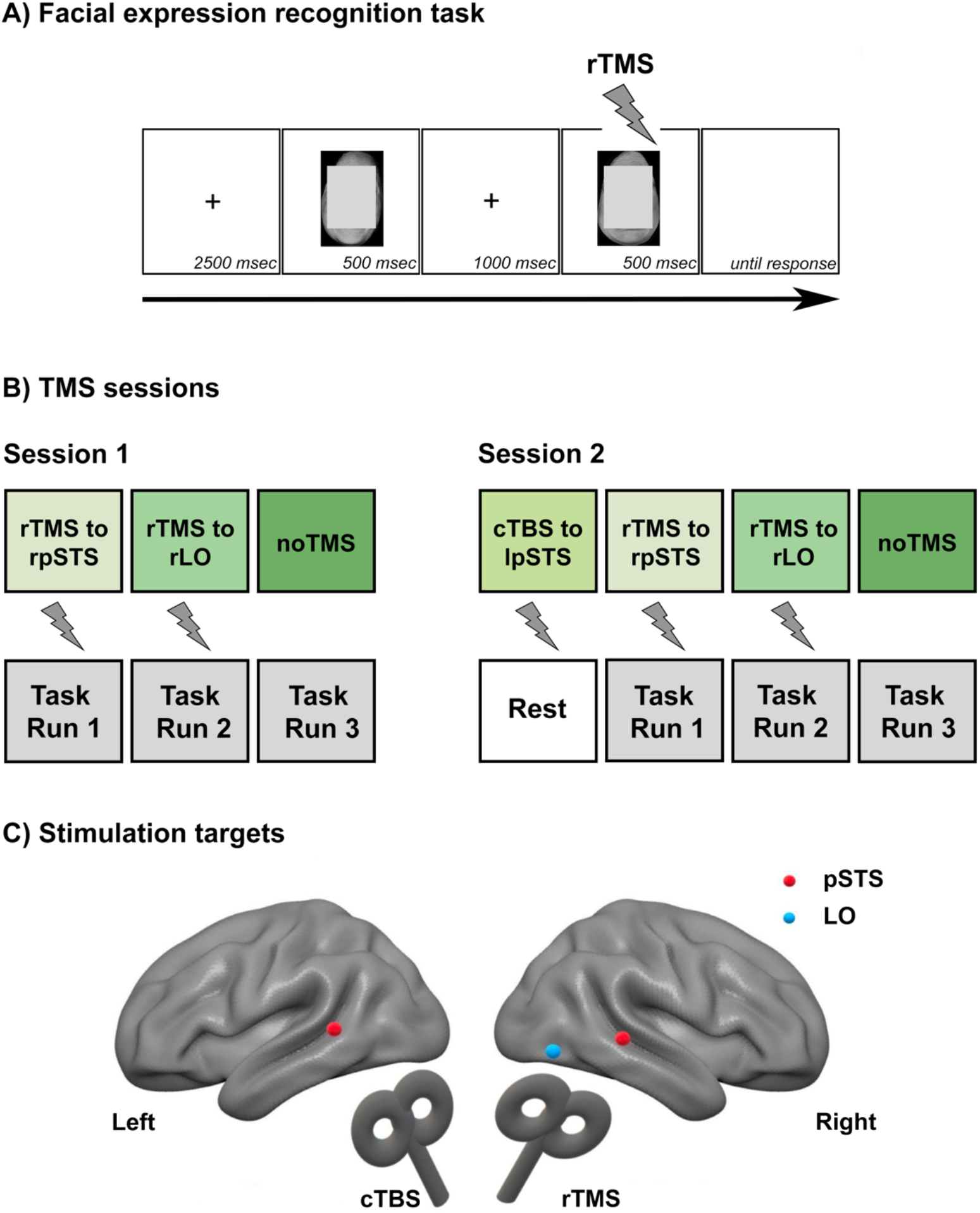
A) An example of a single trial in the facial expression recognition task. B) Experimental procedures during two TMS sessions. During each session, participants performed three runs of the facial expression recognition task during which perturbing rTMS was delivered to either rpSTS (Run 1), rLO (Run 2), or no rTMS was delivered (Run 3). During Session 2 only, conditioning cTBS was delivered to the lpSTS immediately before the task began. C) Illustration of the stimulation targets presented on the standard MNI brain. Note, that stimulation was delivered to targets identified individually for each participant.

During each session, participants completed three runs of the task (see Figure 1B). Each run consisted of 72 trials and lasted approximately 7 minutes. Half of the trials presented the same expressions, while the other half presented different expressions. Each run consisted of the same trials which were presented in a randomised order across runs. Runs were completed under three different stimulation conditions. Perturbing rTMS was delivered on each trial to either the rpSTS (stimulation condition 1) or the rLO (stimulation condition 2). One of the runs acted as a pure behavioural control during which no rTMS was delivered (stimulation condition 3). Stimulation to the object-sensitive site in the rLO was included as a control condition for non-specific effects of rTMS (e.g., facial muscle twitching). This site was chosen as it was not expected to provide any significant effects of stimulation during the expression recognition task. It also has a similar somatosensory sensation of rTMS to the rpSTS as these sites are located in close proximity. The order of the stimulation conditions was randomised across participants but kept the same across two sessions. In Session 2 only, conditioning cTBS was used immediately before the task began.

For on-line perturbing rTMS, we used a train of five pulses delivered at a frequency of 10 Hz (i.e., a pulse every 100 msec) for a duration of 500 msec at a fixed intensity of 60% of the maximum stimulator output. The fixed intensity value was used based on our previous studies (10, 20, 22, 23). Stimulation started at the onset of the second image to maximise a disruptive effect on the expression recognition task. For off-line conditioning cTBS, we used its modified version (24). This stimulation involved a continuous train of 600 pulses delivered in bursts of 3 pulses (a total of 200 bursts) at a frequency of 30 Hz with a burst frequency of 6 Hz for a duration of 33.3 sec and fixed intensity of 50% of the maximum stimulator output. The aim of using cTBS before testing was to induce a longer lasting post-stimulation disruption effect in the lpSTS on the expression recognition task. Goldsworthy and colleagues (14) demonstrated that the effects of the modified cTBS can last up to 30 minutes post-stimulation which would comfortably cover the time of our three task runs. The results of our independent pilot data supported this length of the cTBS effect in pSTS during the expression recognition task. The modified cTBS was used over the standard cTBS as it was shown to produce immediate, longer-lasting, and more reliable effects in contrast to the standard cTBS (14). The order of the sessions with and without cTBS was counterbalanced across participants.

Participants were seated approximately 60 cm away from the computer screen and they provided their responses by pressing the appropriate buttons on a keyboard, using the right index (“same”) or middle (“different”) finger. Participants were instructed to respond as quickly but also as accurately as possible. All stimuli were presented in the centre of a white screen on a Mitsubishi Diamond Pro 2070SB 22-inch CRT monitor, set at 1024 × 768 resolution and refresh rate of 85 Hz. Stimulus presentation and response recording were obtained using E-Prime software (Psychology Software Tools).

#### 2.4.3 Data collection

TMS was delivered using a Magstim Rapid2 stimulator and a Magstim coated Alpha Flat 50 mm diameter figure-of-eight coil (Magstim, Carmarthenshire, UK). The stimulation parameters were within established international safety limits (25, 26). The TMS coil was held against the participant’s head by the experimenter who manually controlled its position throughout testing. All stimulation target sites were marked on each participant’s structural scan using the Brainsight frameless stereotaxy system (Rogue Research, Montreal, Canada). During testing, a Polaris Vicra infrared camera (Northern Digital, Waterloo, ON, Canada) was used in conjunction with the Brainsight to register the participant’s head to their structural scan for accurate stimulation targeting throughout the experiment. All participants wore earplugs in both ears to attenuate the sound of the coil discharge and avoid damage to the ear (27). In some participants, stimulation affected the peripheral jaw muscle and produced a small jaw twitch. Only two participants found TMS over pSTS uncomfortable. Those participants were excluded from the study and no TMS data were collected. The remaining participants tolerated TMS well.

#### 2.4.3 Data analysis

Performance accuracy and reaction times (RTs) were analysed using IBM SPSS Statistics (v24.0) in a 2 × 3 repeated measures ANOVA, with Session (Session 1 with conditioning cTBS to lpSTS and Session 2 without conditioning cTBS to lpSTS) and Stimulation (perturbing rTMS to rpSTS, perturbing rTMS to rLO, and no TMS) as independent factors. Post hoc paired two-tailed t-tests (with Bonferroni correction for multiple comparisons) were used to further characterize significant main effects and interactions from the ANOVA.

## 3. Results

### 3.1 Individual fMRI functional localisation

Face-sensitive areas in the right and left pSTS and object-sensitive area in the rLO were identified in every participant. The peak activations for these areas considerably varied across individuals, matching our previous study (10). The group mean peak coordinates of the stimulation targets in the standard MNI space (rpSTS: *x* = 55, *y* = −38, *z* = 4; lpSTS: *x* = −55, *y* = −43, *z* = 5; rLO: *x* = 50, *y* = −66, *z* = −7) were consistent with our previous studies (10, 20, 22) and studies of others (4, 28, 29).

### 3.2 TMS study

The group mean performance accuracy results are presented in Figure 2. The main effects of Session (F(1, 19) = 10.57; p = 0.004; partial ɲ^2^ = 0.36) and Stimulation (F(2, 38) = 22.38; p < 0.001; partial ɲ^2^ = 0.54) were significant. In addition, a two-way interaction between Session and Stimulation was significant (F(2, 38) = 4.44; p = 0.02; partial ɲ^2^ = 0.19). *Post-hoc t*-tests revealed that perturbing rTMS to the rpSTS during the task significantly impaired expression recognition in both sessions. In Session 1 (without conditioning cTBS to the lpSTS), perturbing rTMS to the rpSTS (83%) produced significantly smaller accuracy than perturbing rpSTS to the rLO (87%; t(19) = 2.96; p = 0.008; *d* = 0.72) and no TMS condition (87%; t(19) = 3.76; p = 0.001; *d* = 0.72). In this session, the impairment effect of perturbing rTMS was recorded in 16 (out of 20) participants. In Session 2 (with conditioning cTBS to the lpSTS), perturbing rTMS to the rpSTS (77%) produced significantly smaller accuracy than perturbing rTMS to the rLO (85%; t(19) = 6.17; p < 0.001; *d* = 1.32) and no TMS condition (85%; t(19) = 5.11; p < 0.001; *d* = 1.23). In this session, the impairment effect of perturbing rTMS was recorded in 17 (out of 20) participants. Crucially in Session 2, perturbing rTMS produced significantly smaller accuracy than in Session 1 (77% vs. 83%; t(19) = 3.34; p = 0.002; *d* = 0.92). This increased impairment effect was present in 15 (out of 20) participants. For the remaining 2 participants, the accuracy was the same across two sessions while for another 3 participants accuracy improved following conditioning cTBS to the lpSTS. The difference between performance accuracy in the control conditions, namely perturbing rTMS to rLO and no TMS, was not different in any of the sessions (both *t*-tests: t(19) < 0.35; p > 0.73; *d* = 0.00). Interestingly, accuracy in the no TMS condition was smaller in Session 2 (85%) than in Session 1 (87%). Although this difference was present numerically, it did not reach the established significance point (t(19) = 1.78; p = 0.09; *d* = 0.36). This indicates, however, that conditioning cTBS to the lpSTS on its own has some effect on expression recognition and this effect is not significantly different from accuracy when only perturbing rTMS was delivered to the rpSTS (83%; t(19) = 1.47; p = 0.16; *d* = 0.33). Nevertheless, both of those single-hemispheric stimulations provided significantly smaller effect on the task than the bilateral (perturbing rTMS to the rpSTS + conditioning cTBS to the lpSTS) stimulation of pSTS.

**Figure 2:**
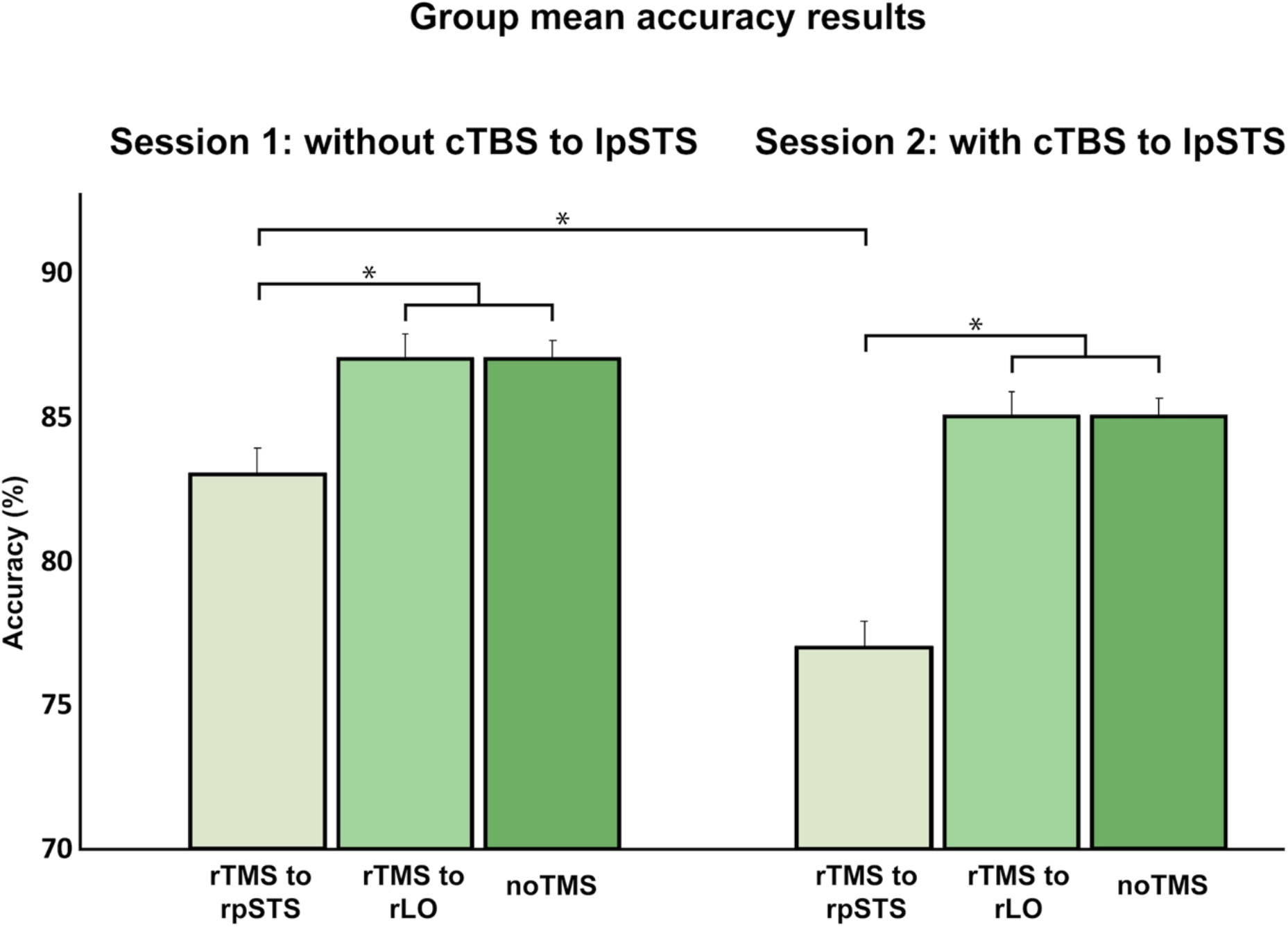
Group mean accuracy results for the facial recognition task under three different stimulation conditions (1: perturbing rTMS to rpSTS, 2: perturbing rTMS to rLO, and 3: no TMS) obtained for the session without (Session 1) and with (Session 2) conditioning cTBS delivered to the lpSTS immediately before the task. Error bars represent *SEM* corrected for repeated measures. \**p* < 0.05.

RTs showed no significant main effects of Session (F(1, 19) = 1.89; p = 0.19; partial ɲ^2^ = 0.90) or Stimulation (F(2, 38) = 3.23; p = 0.05; partial ɲ^2^ = 0.15) and there was no significant twoway interaction between these two conditions (F(2, 38) = 0.53; p = 0.60; partial ɲ^2^ = 0.03).

## 4. Discussion

The current study used a dual-site condition-and-perturb TMS to demonstrate that the lpSTS has a causal functional connection with the rpSTS during facial expression recognition. The conditioning cTBS to the lpSTS, delivered immediately prior to participants performing the facial expression recognition task, doubled the impairment effect of perturbing rTMS to the rpSTS delivered during the task. Importantly, the effect induced by conditioning cTBS was specific to the face network as conditioning cTBS did not affect rTMS to the rLO. This finding causally demonstrates that accurate expression recognition requires functional collaborations between the left and right pSTS.

This study provides further evidence for the importance of the non-dominant hemisphere in cognitive computations and supports face processing models that also include the lpSTS in the face network (e.g., 1). In our prior study (10), we used TMS to demonstrate the casual contribution of the non-dominant pSTS in expression recognition. In our current study, we used TMS to extend this previous finding and demonstrate that the lpSTS is functionally connected to the dominant rpSTS when recognising expressions. Consequently, disruption of the lpSTS combined with disruption of the rpSTS leads to a greater impairment than disruption of these sites separately.

Dual-site condition-and-perturb TMS paradigms offer a unique method to establish the causal functional connectivity between bilateral regions. The only other way to investigate causal functional connectivity between two regions is in lesion studies, which can indicate causal links between brain regions and cognitive functions. However, to our knowledge, no report of a patient with a lesion to both left and right pSTS exists. In addition, the unilateral lesions to the lpSTS that have been reported (e.g., 30, 31, 32) cannot test functional connections of the lpSTS with the rpSTS as there is the possibility of long-term functional re-organisation (33). Over time, the function of a damaged region may be taken over, to various degrees, by other region(s) of the functional network, including the healthy homologue region in the opposite hemisphere. By using dual-site TMS approach, we can simultaneously and temporarily impair a region bilaterally and induce a short-lasting functional impairment, avoiding the issue of anatomical re-organisation (34–36).

Although long-term re-organisation of a cognitive network is not possible using our stimulation protocol, there is a possibility of a short-term re-organisation caused by the conditioning TMS (see, 13). Using fMRI, O’Shea and colleagues demonstrated that conditioning TMS to the left dorsal premotor cortex (PMd), the dominant brain region for action selection of the contralateral hand, resulted in increased activation of regions in the action selection network, including the non-dominant right PMd. After conditioning and perturbing TMS to the left PMd and right PMd, respectively, the action selection was impaired, suggesting the activation increase in the right PMd resulted from functional compensation. As we did not measure the effect of conditioning cTBS on the neural activity in the left or right pSTS, it is impossible to state to what extent our results can be explained by the short-term functional re-organisation. This could be the case if the increased effect of perturbing rTMS to rpSTS results from increased activation in the rpSTS compensating for the lpSTS disfunction. However, the size of the behavioural impairment caused by the condition-and-perturb TMS was greater than individual effects of the conditioning cTBS to the lpSTS or perturbing rTMS to the rpSTS which would indicate that our results are better explained by strong functional connections between the left and right pSTS than only short-term compensation. Future investigations of our effects with fMRI or fNIRS (functional near-infrared spectroscopy) would be of great value for a more detailed understanding of the face network functioning.

## 5. Conclusions

This study provides causal evidence for interhemispheric causal interaction between the left and right pSTS during facial expression recognition. This finding is crucial for our comprehensive understanding of the mechanisms that govern the face network in the human brain and stresses the importance of the non-dominant side of this network. It also advocates dual-site condition-and-perturb TMS as a powerful tool in establishing functional connections between brain regions.

## Acknowledgements

We would like to thank Gesa Hartwigsen for her advice on the participant safety during the dual-site condition-and-perturb TMS.

